# Reliability and validity of multi-band multi-echo fMRI

**DOI:** 10.64898/2026.01.16.699838

**Authors:** Li-Xia Yuan, Ziyang Chen, Chao Jiang, Xiao Chen, Bing-Chen Shao, Zhu-Qing Gong, YiCheng Hsu, Yu-Feng Zang, Xi-Nian Zuo, Hongjian He

**Affiliations:** School of Physics, Zhejiang University, Hangzhou, China; Research Center for Healthcare Data Science, Zhejiang Lab, Hangzhou, China; Beijing Key Laboratory of Learning and Cognition, School of Psychology, Capital Normal University, Beijing, China; Precision TMS Center/Department of Neurology, The Affiliated Hospital of Hangzhou Normal University, Hangzhou, China; Institute of Psychological Sciences, Hangzhou Normal University, Hangzhou, China; Zhejiang Key Laboratory for Research in Assessment of Cognitive Impairments, Hangzhou Normal University, Hangzhou, China; State Key Laboratory of Cognitive Neuroscience and Learning, Beijing Normal University, Beijing, China; MR Research Collaboration Team, Siemens Healthineers Ltd., Shanghai, China; Engineering Research Center of Mobile Health Management System, Ministry of Education, Hangzhou Normal University, Hangzhou, China

**Keywords:** precision neuroimaging, multi-echo fMRI, individual differences, reliability, validity

## Abstract

Precision mapping of individual differences in human brain spontaneous activity is critical for characterizing brain function and guiding clinical translation. Multi-echo resting-state functional MRI (ME-rfMRI) has the potential to provide more reliable mapping of inter-individual differences in intrinsic brain function than conventional single-echo rfMRI (SE-rfMRI). However, the psychometric benefits of ME-rfMRI for mapping individual differences in spontaneous brain activity (SBA) have not been systematically investigated, which is fundamental for understanding the human brain in health and disease. To quantify the psychometric performance of ME-rfMRI in precisely mapping inter-individual variations in SBA, we employed a psychometric design. We scanned 27 healthy adults with both ME-rfMRI and SE-rfMRI to assess short-term (minutes) test-retest reliability, as well as validity across eyes-open (EO) and eyes-closed (EC) resting states. Our results demonstrate that ME-rfMRI improves reliability relative to SE-rfMRI by 10.8% for amplitude of low-frequency fluctuation (ALFF), 4.4% for regional homogeneity (ReHo), and 8.0% for voxel-mirrored homotopic connectivity (VMHC) in cortical regions, and by 12.5% for ALFF, 8.3% for ReHo, and 4.0% for VMHC in subcortical regions. This enhancement is achieved by increasing inter-individual variability while reducing intra-individual variability. Notably, ME-rfMRI accentuates inter-individual variability in the unimodal cortex and clearly delineates two parallel systems with distinct variability patterns within the motor cortex. Concurrently, ME-rfMRI increases effect sizes (4%-8%) for detecting individual differences between EO and EC resting states. Interestingly, ME-rfMRI reveals differential effects of eye closure on two intertwined systems in the primary motor and sensory cortices, extending previous findings. Computational simulations indicate that, compared to SE-rfMRI, ME-rfMRI facilitates experimental designs with reduced costs (i.e., 4%-11% smaller sample sizes) for precise mapping of individual differences. These findings establish, for the first time, the psychometric performance of ME-rfMRI measurements, supporting its potential utility in personalized neuroscience applications, including neurodevelopment and brain disorders.

## INTRODUCTION

Spontaneous brain activity (SBA), which consumes most of the brain’s energy budget for largely unknown functions-termed “dark energy” (Raichle, 2006, 2010, 2015), is highly structured. Resting-state functional magnetic resonance imaging (rfMRI) provides an effective non-invasive method to measure blood-oxygen-level-dependent (BOLD) signals as a proxy for SBA (Biswal et al., 1995; Biswal et al., 2010). Over the three decades since its introduction (Biswal & Uddin, 2025), rfMRI has gained prominence for characterizing brain organization and function, as well as their associations with various behavioral and clinical conditions (Cole et al., 2022; Genon et al., 2022; Gordon et al., 2023; Marek et al., 2022; Zhang & Raichle, 2010).

Recently, mapping multifaceted inter-individual variations in SBA has become a central focus in neuroimaging research (Cui et al., 2020; Dubois & Adolphs, 2016; Genon et al., 2022; Laumann et al., 2023; Mueller et al., 2013; Seghier & Price, 2018; T. Xu et al., 2023; Zilles & Amunts, 2013). Due to the low signal-to-noise ratio (SNR) of conventional single-echo fMRI (SE-fMRI) and limited data acquisition, estimates of inter-individual variability are often contaminated by non-negligible intra-individual variability, particularly in cross-sectional studies (Elliott et al., 2021; C. Jiang et al., 2023; Xing & Zuo, 2018; T. Xu et al., 2023; Zhou et al., 2023; Zuo et al., 2019). Unreliable inter-individual differences impede the discovery of brain-behavior associations and hinder the translation of rfMRI to clinical applications (Brennan et al., 2019; Elliott et al., 2021; Genon et al., 2022; Gratton et al., 2020a; Kong et al., 2021; Noble et al., 2019; Seitzman et al., 2019; Wang et al., 2018; Xing & Zuo, 2018; Zhou et al., 2023).

Multi-echo fMRI (ME-fMRI) has the potential to enhance inter-individual variability while simultaneously reducing intra-individual variability (Elliott et al., 2021; Lynch et al., 2020), thereby providing more reliable mapping of inter-individual differences in intrinsic brain function than SE-fMRI. ME-fMRI acquires multiple whole-brain images at different echo times (TEs) within a single repetition time (TR) (Posse, 2012; Posse et al., 1999), rather than a single volume as in standard SE-fMRI. By optimally combining images acquired at different TEs, BOLD signals can be enhanced (Posse et al., 1999), benefiting the precise characterization of individual traits and their differences. Moreover, ME-fMRI attenuates widely distributed random (thermal) noise (Kang et al., 2023) and denoises SBA signals based on biophysical principles (Kundu et al., 2012, 2013, 2017; Power et al., 2018), substantially minimizing intra-individual variability.

Previous studies have shown that ME-fMRI benefits both task-based experiments and resting-state studies (Gordon et al., 2023; Kang et al., 2023; Lombardo et al., 2016; Lynch et al., 2021, 2020). For task-based experiments, ME-fMRI enhanced effect sizes by up to 149% in mentalizing-associated regions compared to SE-fMRI (Lombardo et al., 2016). For resting-state studies, ME-fMRI (ME-rfMRI) has been shown to increase the within-individual stability of functional connectivity (FC) (Lynch et al., 2021, 2020) and provide superior sensitivity and accuracy for detecting FC and brain networks relative to SE-fMRI (Kang et al., 2023; Kundu et al., 2012, 2013). However, the psychometric benefits of ME-fMRI for mapping individual differences in SBA have not yet been systematically investigated, which is fundamental for understanding the human brain in health and disease (Dubois & Adolphs, 2016; Elliott et al., 2021; Gordon et al., 2023; Raichle, 2015; T. Xu et al., 2023) and crucial for personalized medicine (Gratton et al., 2020b; Laumann et al., 2023; Seghier & Price, 2018; Seitzman et al., 2019; Wang et al., 2018).

To quantify the psychometric performance of ME-rfMRI in precisely mapping inter-individual variations in SBA, we employed a psychometric design. We recruited 27 healthy adults (21 females) and scanned their SBA with both ME-rfMRI and SE-rfMRI to assess short-term (minutes), as well as validity across eyes open (EO and eyes-closed (EC) resting states. Specifically, we investigated inter-individual variability (interV), intra-individual variability (intraV), and reliability of SBA metrics at three spatial scales: amplitude of low-frequency fluctuation (ALFF; Yang et al., 2007a; Zang et al., 2007b), regional homogeneity (ReHo; Jiang & Zuo, 2016; Zang et al., 2004), and voxel-mirrored homotopic connectivity (VMHC; Zuo et al., 2010). We then examined the potential benefits of improved reliability for validity by exploring differences in SBA patterns between two widely used resting conditions (EO and EC) (Yang et al., 2007; L.-X. Yuan et al., 2018; Yue et al., 2023). Furthermore, we performed computational simulations to demonstrate the implications of enhanced psychometric performance on experimental design, examining interplays among reliability, sample size, and effect size (Başol, 2007; Zuo et al., 2013, 2019).

## MATERIALS AND METHODS

### Participants

The study was approved by the Ethics Committee of Zhejiang University (No. 2023-069). All participants provided written informed consent before data acquisition. Participants were screened for any history of neurological or psychiatric disorders. Two MRI datasets were acquired: a reliability dataset and a validity dataset. The reliability dataset included 27 healthy participants (age range: 20-32 years; mean age ± SD: 25 ± 2.9 years; 6 males). The validity dataset included 18 healthy participants (age range: 21-28 years; mean age ± SD: 24 ± 1.6 years; 8 males).

### MRI data acquisition

MRI data were acquired on a Siemens MAGNETOM Prisma 3T scanner (Siemens Healthcare, Erlangen, Germany) with a 64-channel head receive coil. All participants underwent ME-rfMRI, SE-rfMRI, and 3D T1-weighted imaging. Detailed acquisition parameters for each imaging sequence are provided below.

Reliability dataset: T1-weighted images were acquired using a 3D inversion-contrast magnetization-prepared rapid-acquisition gradient echo sequence with a resolution of 1.2 × 1 × 1 mm³. Parameters for SE-rfMRI and ME-rfMRI are listed in Table 1. Data from ME-rfMRI with four different TEs were labeled Echo1, Echo2, Echo3, and Echo4. During resting-state fMRI scanning, participants were instructed to remain still, keep their eyes closed, and avoid focused thinking.

**Table 1.** Imaging parameters for SE-rfMRI and ME-rfMRI in the reliability dataset.

| Imaging Parameter | SE-rfMRI | ME-rfMRI |
| --- | --- | --- |
| Field of view (FOV, mm) |  | 216×216 |
| Matrix size |  | 72×72 |
| Voxel size (mm) |  | 3×3×3 |
| Inter-slice gap (mm) |  | 0 |
| Repetition time (TR, ms) |  | 1300 |
| Flip angle (FA, °) |  | 68 |
| Number of frames per scan |  | 700 |
| Scan duration |  | 15min 10s |
| Multi-band factor | 2 | 3 |
| in plane acceleration factor | 1 | 2 |
| Number of slices | 40 | 42 |
| Echo Spacing (us) | 0.53 | 0.49 |
| Receiver bandwidth (Hz) | 2240 | 2480 |
| Echo time (TE, ms) | 30 | 12.6/30.86/49.13/67.38 |

Validity dataset: T1-weighted images were acquired using a 3D inversion-contrast magnetization-prepared rapid-acquisition gradient echo sequence with a resolution of 0.8 × 0.8 × 0.8 mm³. Each participant underwent four resting-state fMRI sessions, comprising two conditions (EC and EO) and two imaging sequences (ME-rfMRI and SE-rfMRI). The order of the four runs was counterbalanced across participants. Parameters for SE-rfMRI and ME-rfMRI are listed in Table 2. The SE-rfMRI sequence design is similar to that used in landmark fMRI studies, such as the Human Connectome Project (HCP) and the Adolescent Brain Cognitive Development (ABCD) study (Casey et al., 2018; Wall, 2023).

**Table 2.** Imaging parameters for SE-rfMRI and ME-rfMRI in the validity dataset.

| Imaging Parameter | SE-rfMRI | ME-rfMRI |
| --- | --- | --- |
| Matrix size | 90×90 | 72×72 |
| Voxel size (mm) | 2.4×2.4×2.4 | 3×3×3 |
| in plane acceleration factor | 1 | 2 |
| Repetition time (TR, ms) | 750 | 1200 |
| Flip angle (FA, °) | 60 | 68 |
| Number of frames per scan | 600 | 375 |
| Multi-band factor | 6 | 4 |
| Number of slices | 60 | 48 |
| Echo Spacing (us) | 0.50 | 0.49 |
| Receiver bandwidth (Hz) | 2645 | 2572 |
| Echo time (TE, ms) | 30 | 12.6/30.86/49.13/67.38 |
| Field of view (FOV, mm) |  | 216×216 |
| Inter-slice gap (mm) |  | 0 |
| Scan duration |  | 7.5min |

### Data preprocessing

T1-weighted images were preprocessed using fMRIPrep (https://fmriprep.org/en/stable/) with the default anatomical preprocessing pipeline (Esteban et al., 2019). Steps included: 1) intensity non-uniformity correction with N4BiasFieldCorrection; 2) brain extraction using ANTs; 3) brain surface reconstruction with recon-all; 4) nonlinear registration to standard MNI152 space with antsRegistration; and 5) conversion into HCP surface space using ciftify_recon_all from the ciftify workflow (Dickie et al., 2019).

Preprocessing procedures for SE-rfMRI and ME-rfMRI data were kept as similar as possible (Table 3). SE-rfMRI data were preprocessed as follows: timeseries despiking with 3dDespike, slice time correction using 3dTshift, head motion correction using 3dvolreg, registration to the T1-weighted image with bbregister, normalization to standard MNI152 space using transformations from T1-weighted image preprocessing, mapping volume images to HCP surface space with 91k grayordinates, noise signal estimation with ICA-AROMA (Parkes et al., 2018; Pruim et al., 2015), linear trend removal with Nilearn, confounds regression with Nilearn, and band-pass filtering (0.01-0.1 Hz) with Nilearn. Confounds included white matter signal, cerebrospinal fluid signal, global signal, and noise components estimated with ICA-AROMA.

**Table 3.**
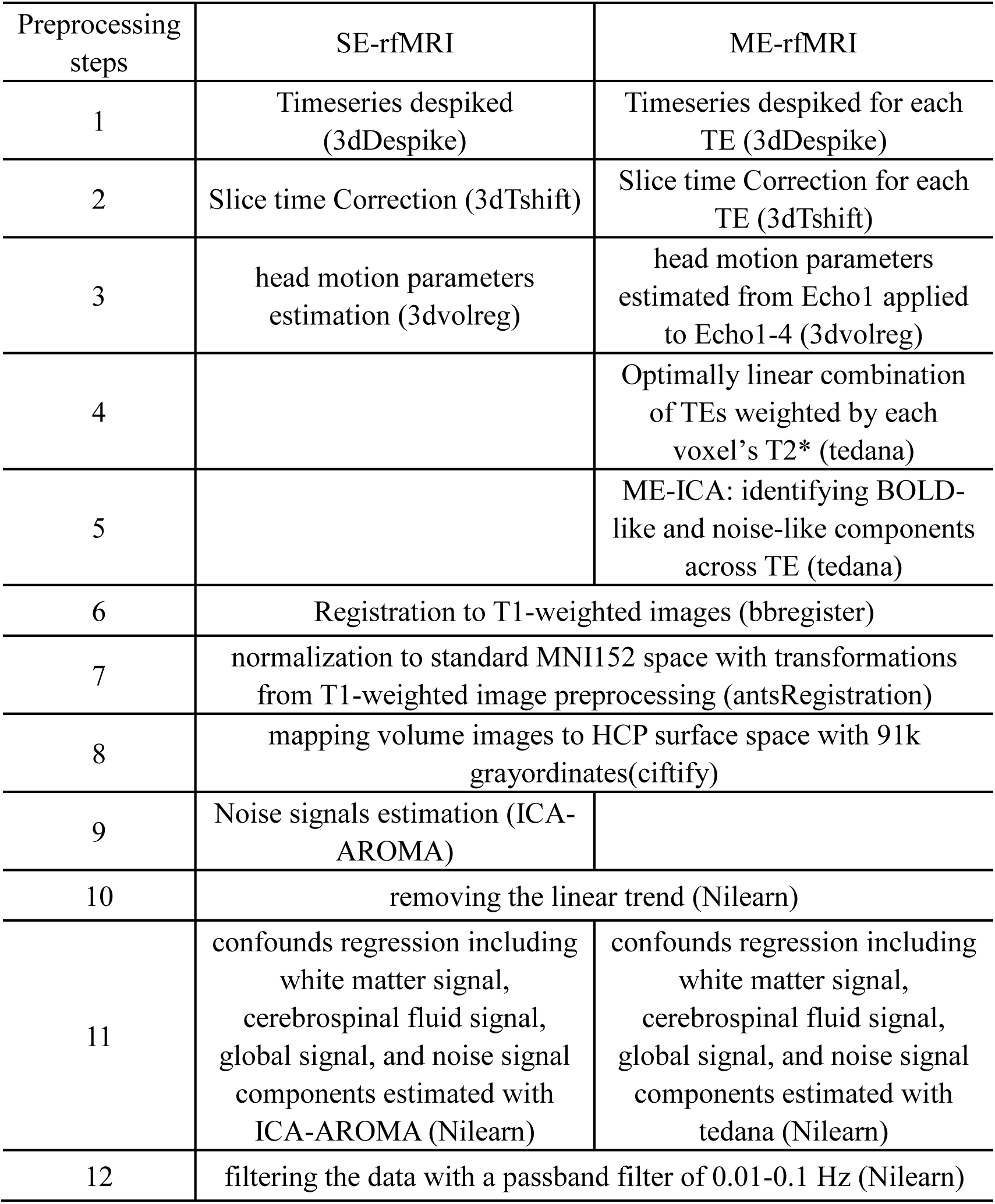
Summary of preprocessing pipelines for SE-rfMRI and ME-rfMRI data.

ME-rfMRI data were preprocessed as follows: timeseries despiking for each echo (Echo1-4), slice time correction for each echo, head motion parameter estimation from Echo1 applied to all echoes, optimal combination of echoes using tedana (DuPre et al., 2021), noise signal estimation using tedana (DuPre et al., 2021; Kundu et al., 2013, 2017), registration to the T1-weighted image, normalization to standard MNI152 space, mapping to HCP surface space, linear trend removal, confounds regression, and band-pass filtering (0.01-0.1 Hz). Confounds included white matter signal, cerebrospinal fluid signal, global signal, and noise signals estimated with tedana.

### Comparison of tSNR

Temporal signal-to-noise ratio (tSNR) was defined as the mean signal divided by the standard deviation of the fMRI time series in MNI space before nuisance regression, using the reliability dataset. Group-level tSNR was obtained by averaging across all subjects within a group mask (retaining voxels present in ≥70% of subjects). tSNR was compared between optimally combined ME-rfMRI, Echo 2 of ME-rfMRI, and SE-rfMRI at both group and subject levels. Two voxels (left orbital frontal cortex [50, 91, 29] and right middle temporal cortex [19, 57, 30]) were selected to illustrate signal fluctuation characteristics, representing challenging and less challenging regions for fMRI.

### Calculation of spontaneous brain activity measurements

After preprocessing, three SBA metrics at different spatial scales (ALFF, ReHo, and VMHC) were computed from both ME-rfMRI and SE-rfMRI time series. ALFF calculation: ALFF measures the mean amplitude of low-frequency fluctuations in the resting-state fMRI time course. ALFF values were calculated as described previously (Yang et al., 2007; Zang et al., 2007). After preprocessing, time series were transformed to the frequency domain using Fast Fourier Transform. The average amplitude within 0.01-0.1 Hz was defined as ALFF. For standardization, ALFF at each vertex was z-transformed across the whole brain to obtain zALFF for further analysis.

ReHo calculation: A surface-based version of ReHo was applied to cortical vertices to reduce partial volume effects and increase test-retest reliability, while a volume-based ReHo was used for subcortical nuclei (L. Jiang & Zuo, 2016; Zuo et al., 2013). ReHo was calculated as Kendall’s coefficient of concordance (KCC) among 6 or 27 nearest neighbors for cortical and subcortical regions, respectively. ReHo values were z-transformed across the whole brain to obtain zReHo.

VMHC calculation: Functional homotopy, the synchrony in spontaneous brain activity between geometrically corresponding interhemispheric (homotopic) regions, is a fundamental trait of intrinsic functional architecture and is measured by VMHC (Zuo et al., 2010). In the symmetrical HCP brain space, Pearson’s correlation coefficients were computed between time series of each vertex and its symmetrical interhemispheric vertex. Correlation values were Fisher Z-transformed and then z-transformed to derive zVMHC.

### Reliability analysis

Short-term test-retest reliability was assessed by splitting the 700-frame rfMRI data from 27 subjects into two sub-datasets of 350 frames each, mimicking two visits.

At the group level, test-retest reliability was characterized by the intra-class correlation coefficient (ICC), which reflects the ratio of inter-individual variability *V_b_* to total variability (sum of inter- and intra-individual variability). As the test-retest rfMRI experiment is a longitudinal design, both intra- and inter-individual variances were estimated using linear mixed models (LMM) (C. Jiang et al., 2023; Zuo et al., 2013). For each vertex, we considered a random sample of *n* subjects with two repeated measurements of a continuous variable *Y* (ALFF, ReHo, or VMHC). Denoting *Y_ij_* as the *i*-th measurement of the *j*-th subject, we applied a two-level LMM to decompose *Y_ij_* as:

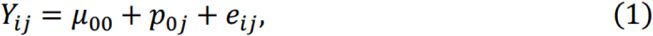

where *μ*_00_ is a fixed parameter, and *p*_0*j*_ and *e_ij_* are independent random effects normally distributed with mean 0 and variances 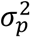 and 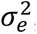, representing inter- and intra-individual variability, respectively. Age, gender, and interval between visits were included as covariates. ICC was defined as:

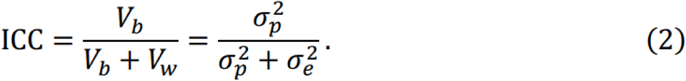

Reliability levels were categorized based on ICC values: slight (0–0.2]), fair ((0.2, 0.4]), moderate ((0.4, 0.6]), substantial ((0.6, 0.8]), and almost perfect ((0.8, 1]) (Landis & Koch, 1977; Noble et al., 2019).

### Validity analysis

Before statistical analysis, spatial smoothing was applied using Gaussian kernels (FWHM = 4 mm for cortical surfaces and subcortical volumes) via Connectome Workbench command line (Marcus et al., 2011). Paired two-tailed *t*-tests were performed to assess SBA differences between EO and EC conditions for zALFF, zReHo, and zVMHC at the vertex level for cortical regions and voxel level for subcortical regions. Significance was set at *p* < 0.05, with further refinement by cluster-size filtering using Connectome Workbench (minimum surface cluster area of 20 mm² for cortical regions, minimum cluster volume of 400 mm³ for subcortical regions).

### Required sample size under different reliabilities of spontaneous brain activity measurements

Interactions between reliability, sample size, and effect size for a statistical power of 0.8 were examined for three experimental designs (two-sample *t*-test, paired *t*-test, and one-way three-level ANOVA) using code shared by Zuo et al. (2019). Required sample sizes at given reliabilities were obtained. The reduction in sample size was calculated as the difference between required sample sizes for ME-rfMRI and SE-rfMRI. The ratio of reduced sample size to required sample size for SE-rfMRI was then computed.

## RESULTS

### tSNR superiority of ME-rfMRI in clinical important regions

Consistent with previous study (Giubergia et al., 2025), optimally combined ME-rfMRI showed substantially higher tSNR (mean tSNR = 71.4) than Echo 2 of ME-rfMRI (mean tSNR = 52.4) (Fig. 1A). Compared to separately acquired SE-rfMRI, ME-rfMRI demonstrated clear advantages in the orbital frontal cortex, inferior temporal cortex, temporal pole, amygdala, hippocampus, brain stem, nucleus accumbens, subgenual anterior cingulate cortex, and cerebellar cortex. Analysis of a representative participant showed that ME-rfMRI better preserved signals in regions with severe signal loss in SE-rfMRI (Fig. 1B), leading to larger effective brain coverage. For example, in the orbital frontal area, a marked increase in mean signal intensity (ME: mean = 6373.7, std = 79.4; SE: mean = 872.5, std = 70.9) combined with comparable temporal signal fluctuation amplitude resulted in much higher tSNR for ME-rfMRI. In contrast, in the middle temporal cortex, despite lower temporal variability in ME-rfMRI (ME: mean = 7526.8, std = 63.6; SE: mean = 12251.1, std = 76.6), substantially lower mean signal intensity led to decreased tSNR compared to SE-rfMRI.

**Fig. 1.**
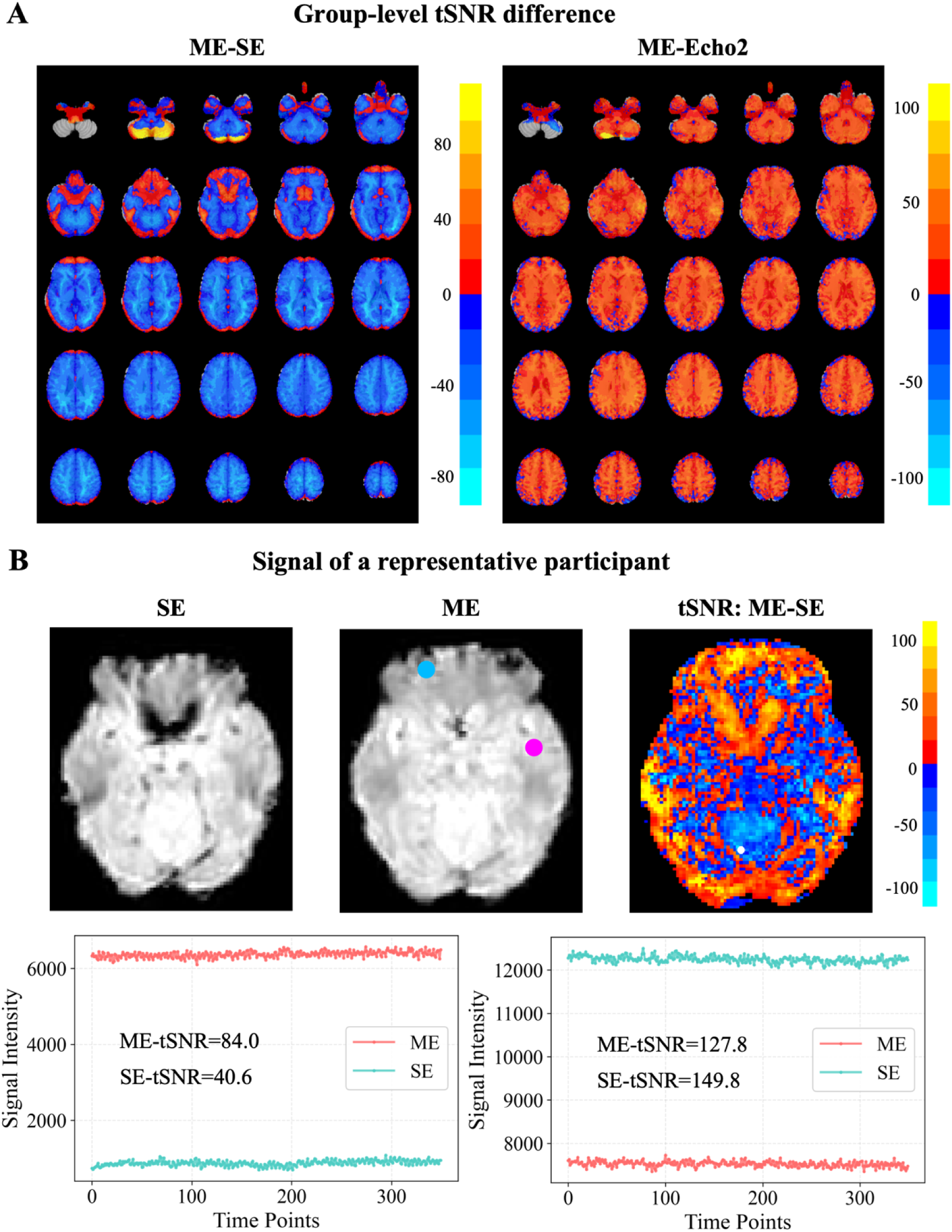
Comparison of temporal signal-to-noise ratio (tSNR) between multi-echo (ME) and single-echo (SE) fMRI acquisitions. (A) Group-averaged voxel-wise tSNR difference maps between optimally combined ME-rfMRI and standard SE-rfMRI data and the second echo from ME-rfMRI. (B) Signal of a representative individual. Top row: mean EPI images and the voxel-wise tSNR difference map (ME minus SE). Cyan and magenta circles indicate the regions of interest (ROIs) used for time-series extraction. Bottom row: raw signal intensity time courses extracted from the corresponding ROIs for SE-fMRI (gray) and ME-fMRI (blue).

### ME-rfMRI achieves highly reliable measures of inter-individual differences

Across the whole brain, ME-rfMRI enhanced inter-individual variability (interV) and reduced intra-individual variability (intraV), resulting in more reliable individual differences in SBA metrics for short-term test-retest designs compared to SE-rfMRI (Fig. 2-3 and Fig. S1-2). The effects of ME-rfMRI exhibited considerable heterogeneity across cortical regions. For instance, more substantial increases in interV and greater reductions in intraV were observed within unimodal cortices, such as the visual and somatomotor networks (Fig. 2, S1, S2). Regarding variability patterns among subnetworks, ME-rfMRI elevated the rank of the somatomotor network and consolidated the high rank of the dorsal attention network (Table S1). Consequently, high interV was observed in the visual, dorsal attention, and default mode networks, while low interV was seen in the limbic and ventral attention networks.

**Fig. 2.**
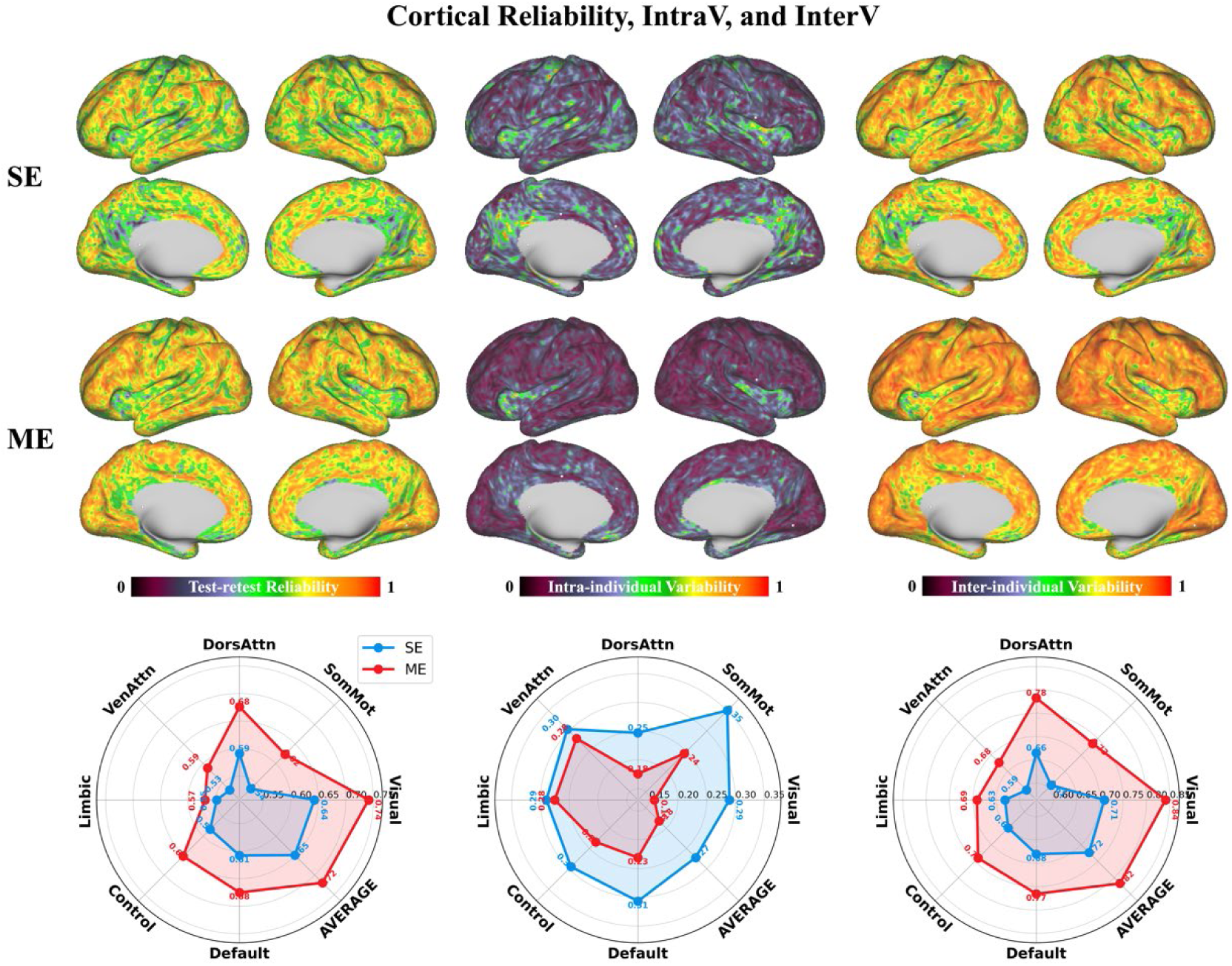
Spatial maps and network profiles of test-retest reliability, intra-individual variability, and inter-individual variability for amplitude of low-frequency fluctuation (ALFF). Results are shown for both multi-echo (ME-rfMRI) and single-echo (SE-rfMRI) acquisitions across the cerebral cortex.

**Fig. 3.**
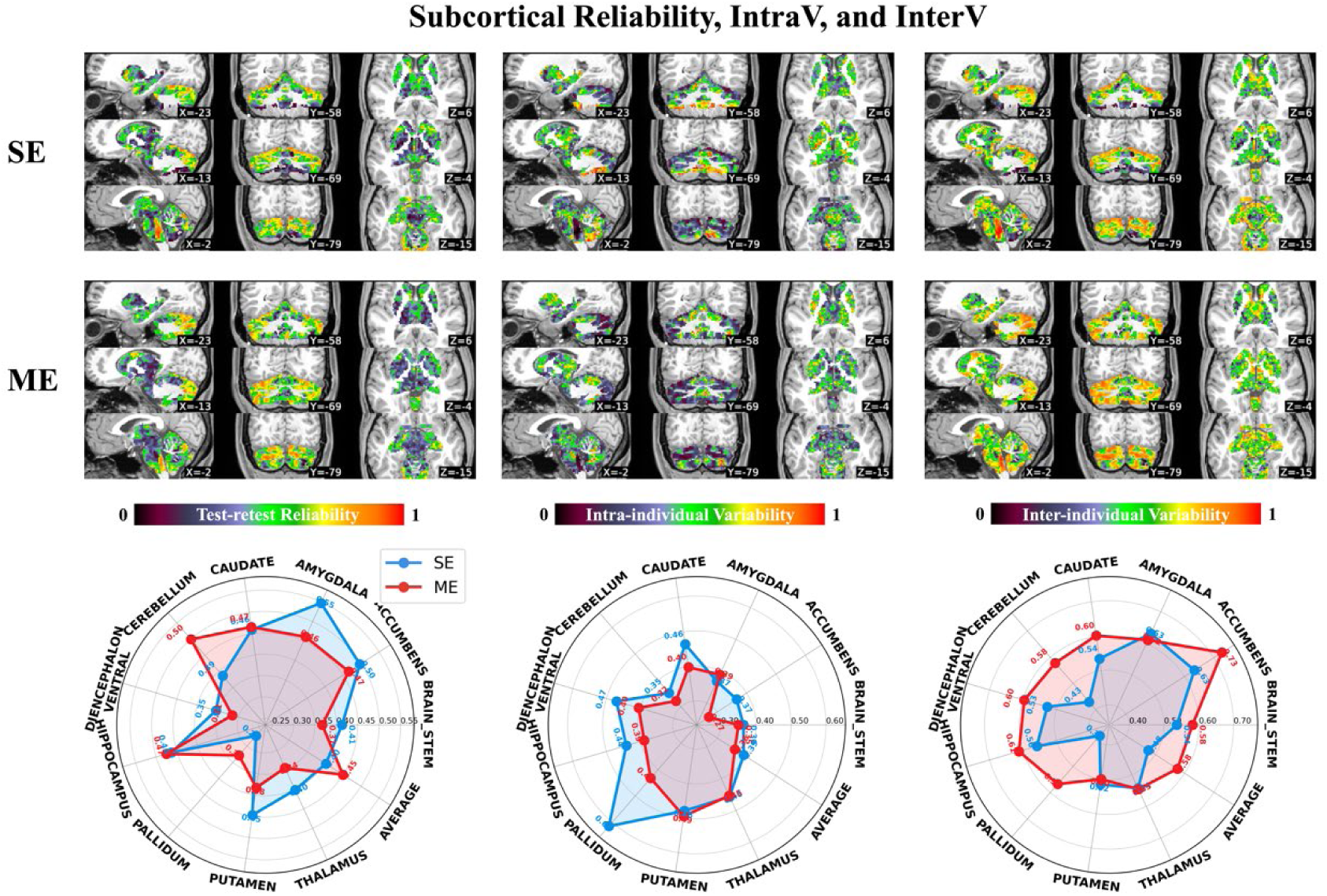
Subcortical maps of test-retest reliability, intra-individual variability, and inter-individual variability for amplitude of low-frequency fluctuation (ALFF). Results are shown for both multi-echo (ME-rfMRI) and single-echo (SE-rfMRI) acquisitions.

Beyond these commonalities, ME-rfMRI effects also varied across the three SBA metrics. ALFF measured with ME-rfMRI exhibited two alternating patterns in the somatomotor cortex on interV and intraV maps (Fig. 2): one with high interV and low intraV, and the other with low interV and high intraV. This echoes recent findings of two parallel systems intertwined within the human motor cortex: effector regions and the somato-cognitive action network (SCAN) (Gordon et al., 2023). ME-rfMRI slightly shifted the interV rank of the default mode network in ALFF (from 2nd to 3rd) while maintaining its rank in ReHo (Table S1). For VMHC, interV rankings remained consistent across methods, with the visual and dorsal attention networks consistently showing the highest variability (Fig. 2 and Fig. S1, S2).

ME-rfMRI generally enhanced reliability across cortical and subcortical regions for all metrics compared to SE-rfMRI (Table S2, S3). Specifically, in the cortex, reliability improvements were 10.8% (ICC from 0.65 to 0.72) for ALFF, 4.4% (ICC from 0.68 to 0.71) for ReHo, and 8.0% (ICC from 0.50 to 0.54) for VMHC. In subcortical regions, improvements were 12.5% (ICC from 0.40 to 0.45) for ALFF, 8.3% (ICC from 0.48 to 0.52) for ReHo, and 4.0% (ICC from 0.25 to 0.26) for VMHC. The visual, dorsal attention, and somatomotor networks benefited most from ME-rfMRI, showing the largest ICC increases (e.g., ALFF increased by 0.10, 0.09, and 0.08, respectively). Among subcortical structures, the cerebellum showed consistent and robust reliability improvements across all metrics (increase of 0.11 for ALFF, 0.08 for ReHo, and 0.07 for VMHC), driving overall subcortical reliability enhancement.

### ME-rfMRI enhances the validity of characterizing individual differences

Compared to the EC condition, ME-rfMRI clearly demonstrated that the EO condition had lower ALFF, ReHo, and VMHC in bilateral sensorimotor and auditory cortices, and higher values in the default mode and control networks (Fig. 4), consistent with previous studies (Feng et al., 2021; Jia et al., 2020; Liu et al., 2013; B. Yuan et al., 2014). Interestingly, the ventrolateral part of the visual network showed lower intrinsic metrics, while the dorsomedial part showed higher metrics under EO. Notably, brain regions with EO vs. EC differences were alternately distributed in three clusters within the primary somatosensory cortex in ME-rfMRI, corresponding to SCAN rather than effector regions (Gordon et al., 2023). This pattern was most evident in ALFF, followed by ReHo, and then VMHC. However, these findings were not apparent in SE-rfMRI.

**Fig. 4.**
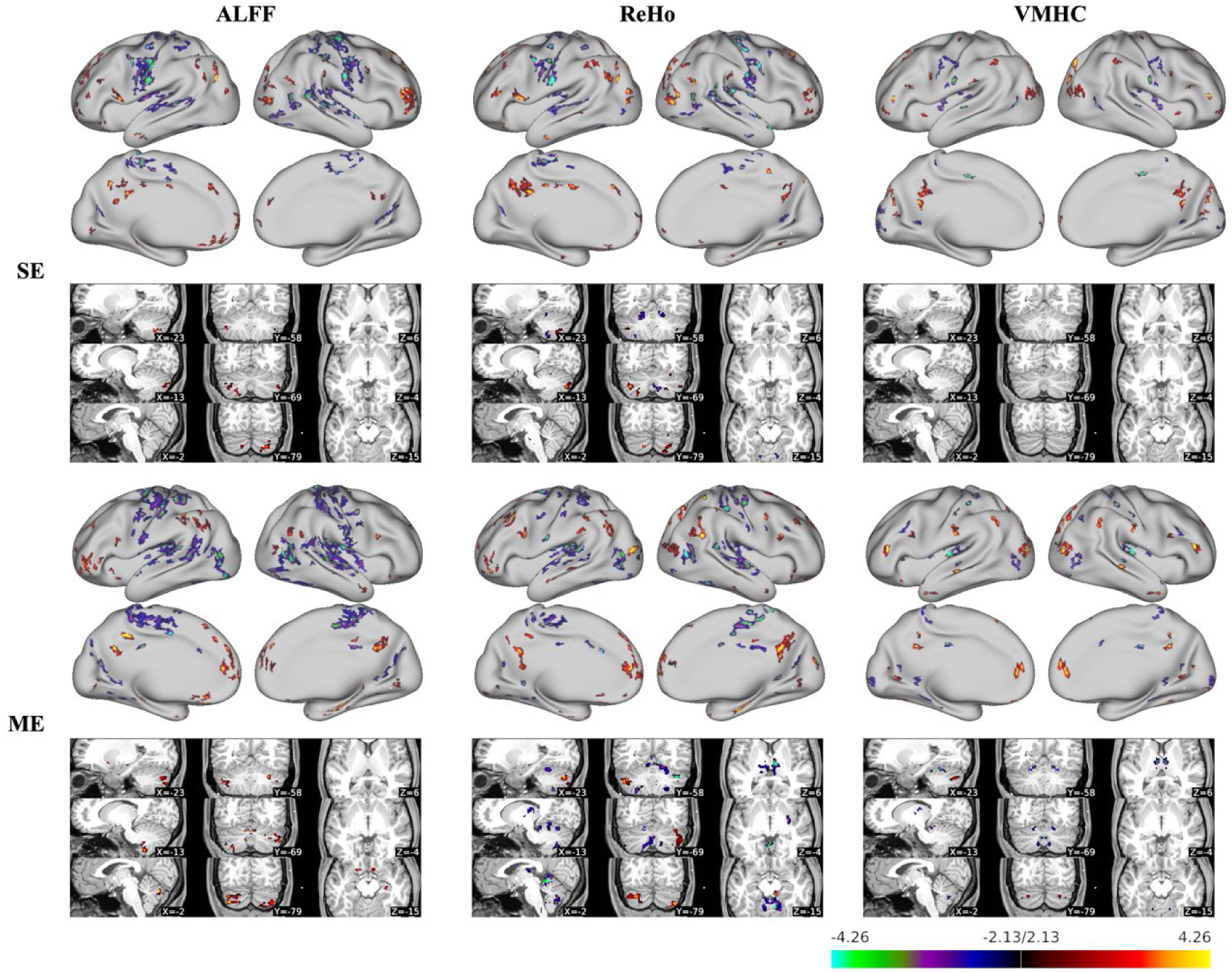
Paired *t*-test maps comparing eyes open (EO) and eyes closed (EC) states. Statistical maps depict differences in ALFF, ReHo, and VMHC for both cortical (surface) and subcortical (volume) regions. Warm colors (red-yellow) indicate EO > EC; cool colors (blue-cyan) indicate EO < EC.

For subcortical structures, ME-fMRI revealed significant EO vs. EC differences in the cerebellum and thalamus that were largely absent or weaker in SE-fMRI. Specifically, the cerebellum showed decreased intrinsic metrics during EO, while the thalamus exhibited increased activity, consistent with their roles in sensorimotor processing and arousal modulation.

Mean effect size increases due to ME-rfMRI were 4.6% for ALFF, 6.0% for ReHo, and 7.3% for VMHC. Effect size increases showed spatial dependence across brain networks. More substantial boosts were observed in the visual network (ALFF: 5.0%, ReHo: 8.5%, VMHC: 7.6%) and default mode network (ALFF: 9.4%, ReHo: 4.7%, VMHC: 6.5%), with consistent improvements across metrics. The somatomotor network also showed notable enhancements, particularly in VMHC (10.3%) and ReHo (5.3%). The dorsal attention network demonstrated marked improvement in ReHo (9.8%).

### ME-rfMRI optimizes the experimental design

Increased reliability can substantially reduce sample size requirements for achieving appropriate statistical power in neuroscience, where effect sizes are often modest to moderate. We quantified the benefit of ME-rfMRI in reducing sample size based on short-term reliability. Our findings indicated that greater sample size reductions occurred for metrics with lower initial reliability and larger reliability enhancements, such as VMHC. For example, with an effect size of 0.2, sample size reductions (and cost savings) in cortical regions were approximately 10.8% for ALFF, 4.4% for ReHo, and 8.0% for VMHC (Fig. 5A). For subcortical regions, corresponding reductions were 12.5% for ALFF, 8.3% for ReHo, and 4.0% for VMHC (Fig. 5B). These reductions remained stable across various experimental designs and effect sizes.

**Fig. 5.**
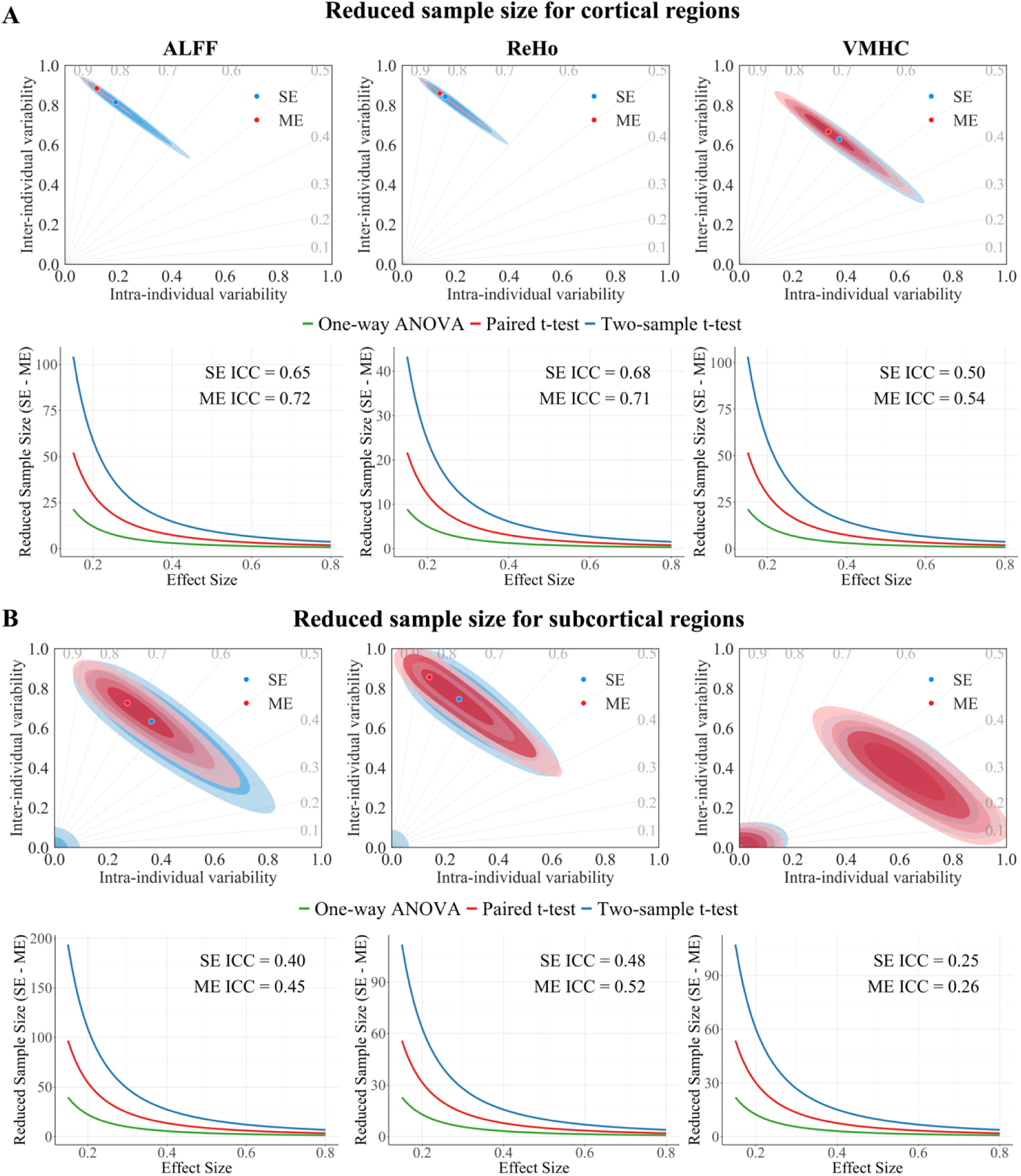
Sample size implications of improved reliability with ME-fMRI. (A, B) For cortical and subcortical regions, respectively, the joint distribution of intra- and inter-individual variability is shown for SE-rfMRI and ME-rfMRI (ALFF, ReHo, VMHC). Corresponding reductions in required sample size are plotted for different effect sizes and experimental designs, calculated based on the change in reliability from SE-rfMRI to ME-rfMRI.

## DISCUSSION

Individual differences in spontaneous brain activity have gained increasing attention due to their relevance for characterizing brain organization and dynamics in both healthy and clinical populations. Precise mapping of individual variability in SBA has been challenging due to the low reliability and validity of conventional SE-rfMRI. In this study, we employed ME-rfMRI to enhance reliability and validity. For the first time, we demonstrate that ME-rfMRI improves reliability by increasing inter-individual variability and reducing intra-individual variability, and boosts validity for detecting individual differences across multiple spatial-scale intrinsic metrics. We also show that ME-rfMRI enhances inter-individual variability in unimodal cortices, highlighting their importance relative to transmodal cortices. Unlike SE-rfMRI, ME-rfMRI facilitates detection of two parallel systems intertwined in the motor cortex in both inter-individual variability maps and brain activity difference maps between EO and EC, indicating unique advantages for precise mapping of individual differences.

### Precision mapping of individual differences in spontaneous brain activity

The fact that individuals think and behave differently stems from individual differences in brain anatomy and connectivity (Mueller et al., 2013). Considering brain function at the individual level enables fine-grained mapping of functional anatomy, determination of whether an individual scan is normative relative to a reference population, and understanding sources of inter-individual variability in brain activity (Van Horn et al., 2008), which is crucial for personalized medicine and understanding neural bases of human cognition and behavior (Kong et al., 2019). RfMRI is a key tool for studying individual differences due to its ease of implementation and validity for depicting brain organization (Biswal et al., 2010; Dubois & Adolphs, 2016). However, BOLD signals from SE-fMRI are susceptible to non-neural confounds, such as cardiac and respiratory physiology, motion, thermal noise, and other sources (Kundu et al., 2012; Power et al., 2018), raising concerns about fMRI reliability in both neuroscience and clinical settings.

For example, functional connectivity, one of the most common rfMRI-derived metrics, exhibits poor test-retest reliability (typically around 0.29) (Noble et al., 2019), far below the desired clinical standard (0.8) (Xing & Zuo, 2018). This inadequate reliability hinders the translation of fMRI to clinical applications and impedes studies of individual differences in brain function (Elliott et al., 2021; Zhao et al., 2022). For the first time, we demonstrate that ME-rfMRI improves reliability by 4.0%–12.5% for multiple spatial-scale intrinsic metrics.

Although ME-rfMRI has been used in various contexts (Alvand et al., 2022; Cohan et al., 2023; Gordon et al., 2023; Hearne et al., 2019), exploration of its benefits for characterizing individual differences has lagged. Our study addresses this gap by providing pioneering evidence of its potential for precise mapping of inter-individual variability in SBA through reduction of confounding intra-individual variability. These findings align with the unique technical features of ME-rfMRI. ME-rfMRI simultaneously acquires multiple images with different TEs and BOLD contrast within one TR, with signals decaying exponentially with TE (Kundu et al., 2017; Yuan et al., 2021). Images from different TEs can be combined based on T2* decay rates at each vertex to create an “optimally combined” time series (Posse et al., 1999), akin to using a customized TE for each vertex. For instance, signal dropout can be recovered in the orbital frontal cortex (with short T2*), and BOLD contrast can be improved in the prefrontal cortex (with long T2*) (Lynch et al., 2020). Thus, ME-fMRI enhances neurological signals for precise delineation of individual traits. Furthermore, combining multiple echoes attenuates thermal (random) noise (Kang et al., 2023), which can constitute a large fraction of recorded signals (Power, 2017). As thermal noise is embedded in all fMRI signals, this effect occurs throughout the brain. Additionally, signal decay across echoes can be used during denoising to identify and remove non-neurobiological signals related to head motion, breathing, hardware instability, and cerebrovascular pulsation (Kundu et al., 2012, 2013, 2017; Power et al., 2018). Thus, ME-rfMRI suppresses noise and non-neurological signals to minimize intra-individual contamination. Collectively, these technical advantages of ME-rfMRI underpin our solid findings of increased inter-individual variability and reduced intra-individual variability in short-term test-retest designs.

### Individual differences: unimodal versus transmodal cortex

Regarding inter-individual variability patterns across metrics, our results show that ME-fMRI markedly elevated the rank of auditory and somatosensory-motor networks, highlighting the importance of exploring individual differences in unimodal cortex. This contrasts with previous whole-brain functional connectivity findings that association cortex is more variable than unimodal cortex (Kong et al., 2019; Mueller et al., 2013). Additionally, our results reveal that inter-individual variability patterns of VMHC resemble its amplitude patterns across the brain (Zuo et al., 2010), suggesting that stronger interhemispheric functional connectivity tends to vary more across individuals. Previous seed-based functional connectivity studies of the posterior cingulate cortex also showed that inter-individual variability patterns overlapped with group-level statistical significance maps (Chen et al., 2015). These findings reflect strength-dependent inter-individual variability patterns of functional connectivity, consistent with reports that functional variability is positively associated with long-range connectivity but negatively associated with local connectivity (Mueller et al., 2013). For example, high long-range connectivity in transmodal cortex indicates stronger whole-brain connections, while high local connectivity in unimodal cortex implies weaker brain-wide connections. Correspondingly, our results show that high intra-individual variability of VMHC is localized in regions with low functional connectivity, leading to low reliability, consistent with previous studies (Chen et al., 2015). Furthermore, local metrics (e.g., ALFF) reflect vertex-specific properties rather than functional connections, exhibiting greater temporal stability than connectivity metrics. Collectively, multiple intrinsic metrics provide a more comprehensive picture of functional individual variability.

Recent studies demonstrate that the classic homunculus in motor cortex comprises both SCAN and effector regions, with these two parallel systems forming an integrate-isolate pattern: effector-specific regions (foot, hand, mouth) for fine motor control isolation, and SCAN for integrating goals, physiology, and body movement (Gordon et al., 2023). Correspondingly, we show that motor cortex exhibits two alternating patterns in individual variability. Based on spatial localization, we infer that SCAN shows higher inter-individual variability than effector regions in ALFF, potentially emphasizing SCAN’s role in human behavior and cognition. ME-fMRI facilitates identification of the intertwined parallel systems in motor cortex, particularly in ALFF compared to ReHo and VMHC, which may be attributed to distinct characteristics of these metrics. ALFF depicts local activity and has the highest reliability (Jia et al., 2020; Yang et al., 2007; Y.-F. Zang et al., 2007). The natural smoothing effect of ReHo may hinder separation of SCAN and effector regions (Yufeng Zang et al., 2004; Zuo et al., 2013). VMHC has moderate reliability, with intra-individual variability dominating inter-individual variability in most brain regions, and its reliance on coordinate symmetry rather than functional symmetry may limit efficiency (Zuo et al., 2010).

### Beyond reliability: the validity of characterizing individual differences

Reliability is a prerequisite for validity and sets an upper bound for validity; however, high reliability can be driven by artifacts rather than meaningful signals (Zuo et al., 2019). Using construct validity through intrinsic functional differences between EO and EC conditions, we further explored potential benefits of improved reliability for validity enhancement with ME-rfMRI. Our findings demonstrate that whole-brain reliability enhancement with ME-fMRI greatly improves validity (i.e., effect sizes). For example, with elevated reliability in ME-rfMRI, decreased ALFF in the default mode network under EC versus EO can be clearly detected, consistent with previous research (Liu et al., 2013; Yang et al., 2007; Yuan et al., 2014; Yue et al., 2023). For sensory-motor and auditory networks, reliability enhancements also substantially boost effect sizes for all spatial metrics. Previous studies showed that ME-fMRI increased effect sizes by up to 149% in mentalizing-related regions compared to SE-fMRI (Lombardo et al., 2016). As a supplement, we demonstrate great benefits of ME-fMRI for improving validity in detecting individual variability in SBA. Consequently, statistical power can be markedly enhanced, facilitating reproducible findings and alleviating the confidence crisis in fMRI (Button et al., 2013).

Volitional eye closure is observed only in conscious, awake humans and is rare in animals. EC and EO are fundamental behaviors for directing attention to the internal milieu versus the external world, representing different states of awareness (Feng et al., 2021; Han et al., 2024; Tang et al., 2015). Neuroimaging studies have established that eye closure enhances brain activity in primary sensory-motor regions (Brodoehl et al., 2015; Liu et al., 2013; P. Xu et al., 2014; Yue et al., 2023). In this study, we detected two alternating patterns of brain function differences between EO and EC in sensory-motor cortex. Interestingly, one pattern showed significantly higher local spontaneous activity, regional homogeneity, and inter-hemispheric coherence under EC versus EO and appeared located in SCAN, while the other showed no significant difference and was distributed in effector regions. This finding aligns with previous research on SCAN function and physiological differences between EO and EC. SCAN is believed to be critical for action planning, physiological control, arousal, error processing, and pain (Gordon et al., 2023). Correspondingly, physiological differences between EO and EC relate to self-awareness, imagination, and arousal states rather than practical somatomotion (Marx et al., 2003, 2004; Yan et al., 2009). However, further task-related investigations are needed for thorough verification.

### Experimental implications

Neuroscience aims to understand how the nervous system relates to behavior and to guide clinical care in neurological and psychiatric disorders (Gratton et al., 2022). Due to low reliability and small effect sizes, one proposed solution for robust brain-behavior correlations across individuals is to collect large, consortia-sized samples (Tiego et al., 2023). For example, Marek et al. (2022) demonstrated that cross-sectional brain-behavior correlations are often small, and reproducible brain-wide association studies require thousands of individuals. However, increasing sample sizes will have limited impact unless a more fundamental issue is addressed: the reliability of measurements used to depict spontaneous brain activity. By increasing reliability, required sample sizes are dramatically reduced for achieving appropriate statistical power in neuroscience, where effect sizes range from modest to moderate (Zuo et al., 2019), as fully reflected in our results (Figure 5). Moreover, our results show that sample size reduction with ME-rfMRI is 4.0%-12.5% across different experimental designs and effect sizes compared to SE-fMRI, facilitating cost and labor savings. Importantly, enhanced reliability with ME-fMRI also suggests potential for improving efficiency in individual-level precision mapping, which typically requires long scan durations (Lynch et al., 2020). By yielding more robust measurements, ME-fMRI could improve clinical feasibility for populations unable to tolerate extensive scanning, such as children and psychiatric patients.

### Limitations

This study has certain limitations. First, scan duration for short-term test-retest reliability was 7.5 minutes, which may be insufficient for obtaining highly reliable VMHC. Longer scan durations are needed to further explore reliability variations with imaging time for VMHC. Second, while optimization of acquisition protocols (e.g., selection of optimal TR, TE, flip angle, acceleration level, spatial resolution) and data processing are beyond the scope of this study, they remain important and active research areas for ME-rfMRI.

### Conclusions

Our study demonstrates that ME-rfMRI significantly improves reliability for intrinsic SBA metrics and enhances validity for detecting individual differences. Furthermore, ME-rfMRI accentuates inter-individual variability in unimodal cortices and precisely delineates two parallel systems intertwined in motor cortex. Additionally, ME-rfMRI facilitates reductions in sample size, experimental costs, and scan durations for precise individual difference mapping. Together, these findings establish the potential utility of ME-rfMRI for precise inter-individual variability mapping and suggest a compelling future role for ME-rfMRI in both neuroscience and

## DATA AND CODE AVAILABILITY

The data and code that support the findings of this study are available from the corresponding author, H.H., upon reasonable request. The data and code are not publicly available due to privacy or ethical restrictions.

## AUTHOR CONTRIBUTIONS

L.Y.: Data curation, Formal analysis, Methodology, Software, Writing - original draft, Writing - review & editing. Z.C: Data curation, Formal analysis, Software, Writing - original draft. J.C.: Data curation, Formal analysis, Methodology, Software, Writing - original draft. X.C.: Formal analysis, Software,Visualization. B.S. and Z.G.: Formal analysis, Software. Y.H.: Data curation, Software. Y.Z.: Methodology, Writing-review & editing, Funding acquisition. X.Z.: Conceptualization, Formal analysis, Methodology, Writing-review & editing. J.H.: Conceptualization, Methodology, Formal analysis, Writing-review & editing, Supervision.

## ACKNOWLEDGEMENTS

We thank Qiuping Ding from the Center for Brain Imaging Science and Technology, Zhejiang University, for her technical assistance with the MRI scanning.

## SUPPLEMENTARY MATERIALS

**Fig. S1.**
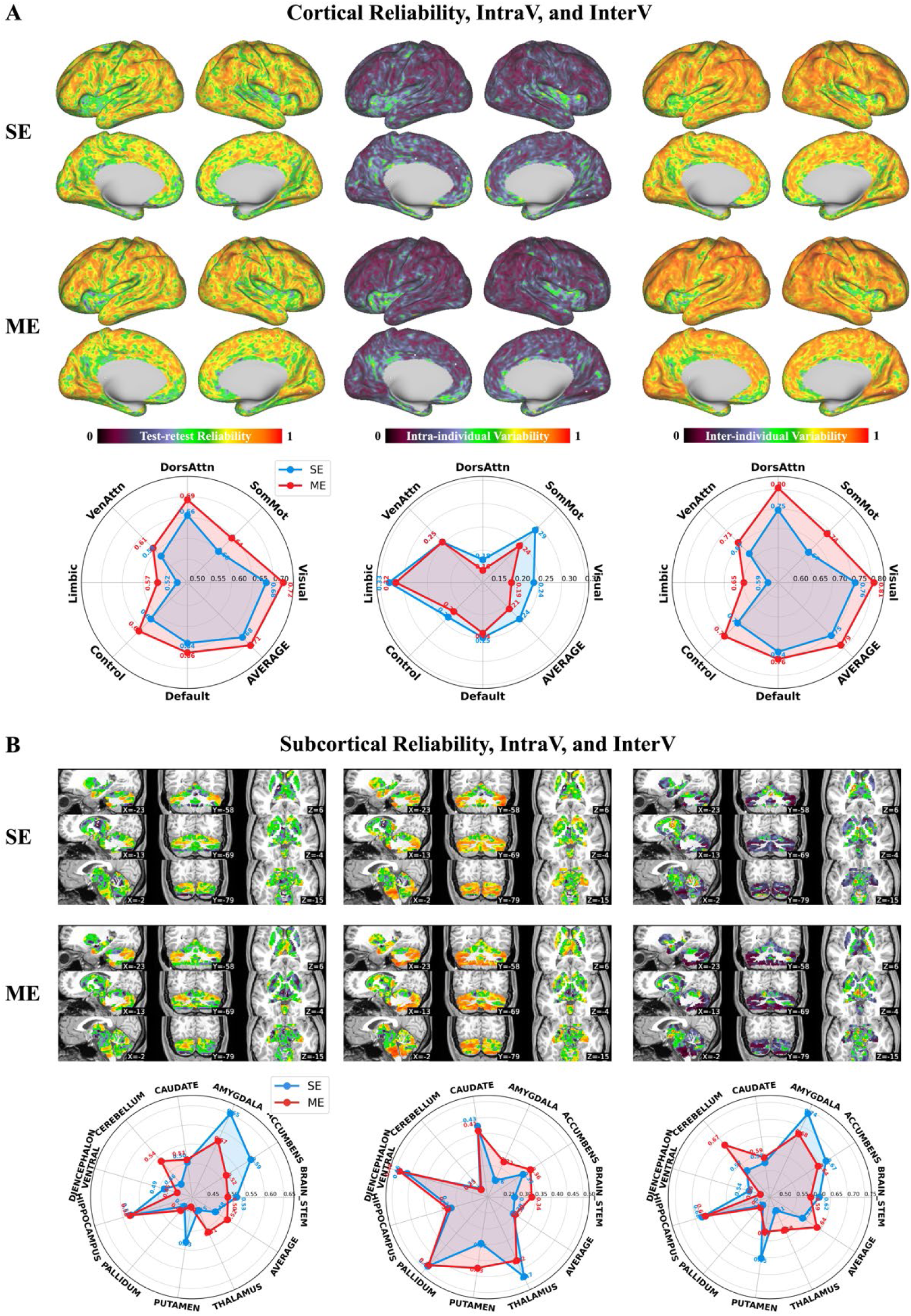
Reliability, intra- and inter-individual variability of regional homogeneity (ReHo). (A) Spatial distribution and network profiles across the cerebral cortex for ME-fMRI and SE-rMRI. (B) Corresponding maps and values for subcortical regions.

**Fig. S2.**
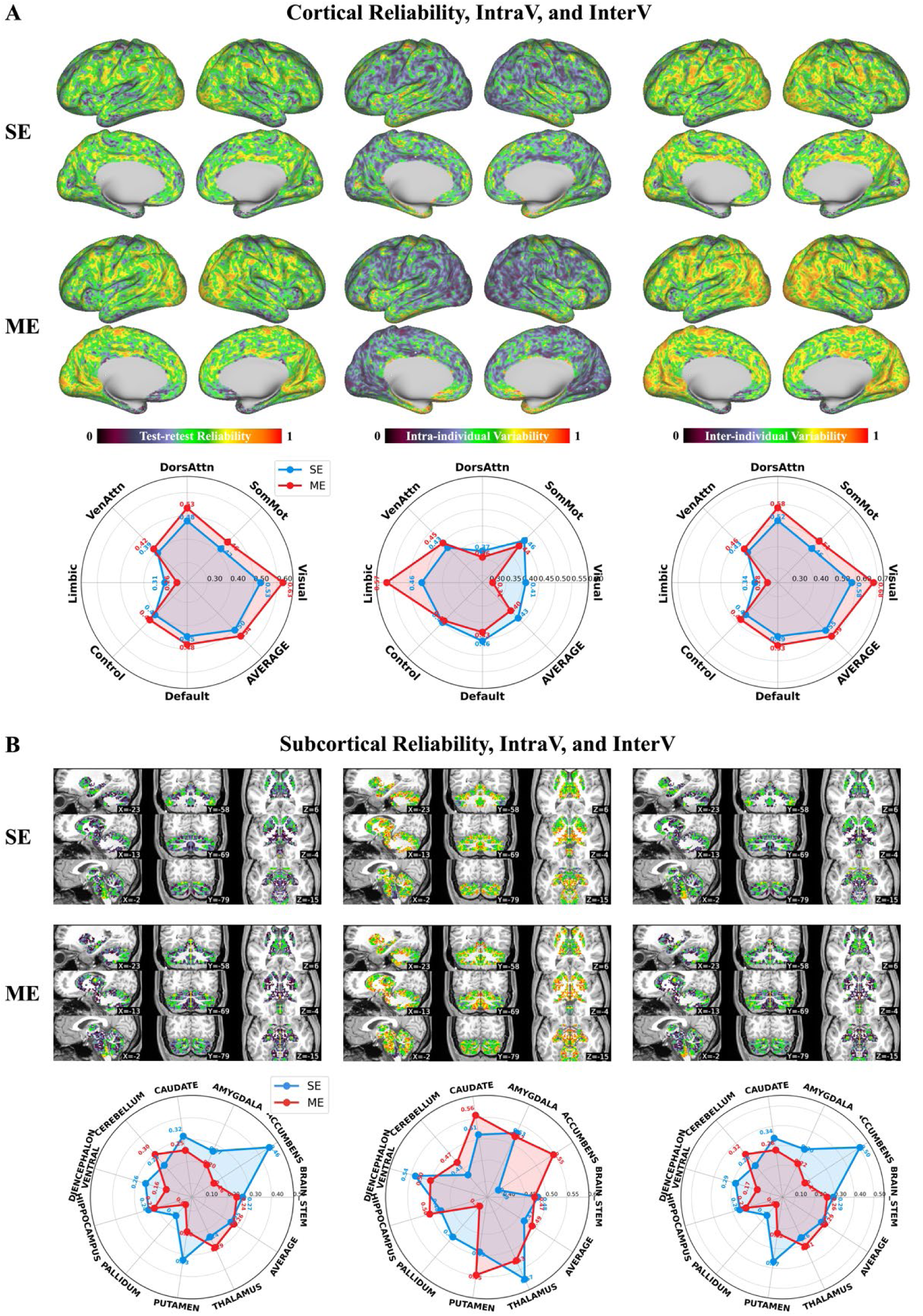
Reliability, intra- and inter-individual variability of voxel-mirrored homotopic connectivity (VMHC). (A) Spatial distribution and network profiles across the cerebral cortex for ME-fMRI and SE-fMRI. (B) Corresponding maps and values for subcortical regions.

**Table S1.** Mean values and rankings of inter- and intra-individual variability across seven cortical networks. Data are shown for both ME-fMRI and SE-fMRI.

|  | Metrics | SomMot | Visual | VenAttn | DorsAttn | Control | Default | Limbic |
| --- | --- | --- | --- | --- | --- | --- | --- | --- |
| InterV<br>(mean/rank) | ME-ALFF | 0.73/5 | 0.84/1 | 0.68/7 | 0.78/2 | 0.74/4 | 0.77/3 | 0.69/6 |
|  | SE-ALFF | 0.61/6 | 0.71/1 | 0.59/7 | 0.66/3 | 0.65/4 | 0.68/2 | 0.63/5 |
|  | ME-ReHo | 0.74/4 | 0.81/1 | 0.71/5 | 0.80/2 | 0.76/3 | 0.76/3 | 0.65/6 |
|  | SE-ReHo | 0.67/5 | 0.76/1 | 0.67/5 | 0.75/2 | 0.71/4 | 0.74/3 | 0.59/6 |
|  | ME-VMHC | 0.51/4 | 0.68/1 | 0.46/6 | 0.58/2 | 0.47/5 | 0.53/3 | 0.28/7 |
|  | SE-VMHC | 0.46/4 | 0.58/1 | 0.43/6 | 0.52/2 | 0.45/5 | 0.49/3 | 0.34/7 |
| IntraV<br>(mean/rank) | ME-ALFF | 0.24/3 | 0.16/7 | 0.28/1 | 0.18/6 | 0.23/4 | 0.23/4 | 0.28/1 |
|  | SE-ALFF | 0.35/1 | 0.29/5 | 0.30/3 | 0.25/7 | 0.30/3 | 0.31/2 | 0.29/5 |
|  | ME-ReHo | 0.24/3 | 0.19/6 | 0.25/2 | 0.16/7 | 0.22/5 | 0.24/3 | 0.32/1 |
|  | SE-ReHo | 0.29/2 | 0.24/5 | 0.25/3 | 0.18/7 | 0.24/5 | 0.25/3 | 0.33/1 |
|  | ME-VMHC | 0.46/2 | 0.31/7 | 0.45/3 | 0.37/6 | 0.44/4 | 0.43/5 | 0.57/1 |
|  | SE-VMHC | 0.44/4 | 0.41/6 | 0.43/5 | 0.36/7 | 0.45/3 | 0.46/1 | 0.46/1 |

**Table S2.** Test-retest reliability (ICC) of intrinsic metrics (ALFF, ReHo, VMHC) across cortical networks. Reliability values are compared between ME-fMRI and SE-fMRI acquisitions.

|  | Metrics | DorsAttn | SomMot | Visual | Default | Control | Limbic | VenAttn | Average |
| --- | --- | --- | --- | --- | --- | --- | --- | --- | --- |
| Cortical<br>ICC<br>(ME/SE) | ALFF | 0.68/0.59 | 0.62/0.54 | 0.74/0.64 | 0.68/0.61 | 0.65/0.58 | 0.57/0.55 | 0.59/0.53 | 0.72/0.65 |
|  | ReHo | 0.69/0.66 | 0.64/0.60 | 0.72/0.68 | 0.66/0.64 | 0.66/0.62 | 0.57/0.52 | 0.61/0.58 | 0.71/0.68 |
|  | VMHC | 0.69/0.66 | 0.64/0.60 | 0.72/0.68 | 0.66/0.64 | 0.66/0.62 | 0.57/0.52 | 0.61/0.58 | 0.54/0.50 |

**Table S3.** Test-retest reliability (ICC) of intrinsic metrics (ALFF, ReHo, VMHC) across subcortical regions. Reliability values are compared between ME-fMRI and SE-fMRI acquisitions.

| Metric | Brainstem | Accumbens | Amygdala | Caudate | Cerebellum | Diencephalon_v | Hippocampus | Pallidum | Putamen | Thalamus | Average |
| --- | --- | --- | --- | --- | --- | --- | --- | --- | --- | --- | --- |
| ALFF | 0.37/0.41 | 0.47/0.50 | 0.46/0.55 | 0.47/0.46 | 0.50/0.39 | 0.31/0.35 | 0.47/0.46 | 0.33/0.27 | 0.38/0.48 | 0.34/0.40 | 0.45/0.40 |
| ReHo | 0.50/0.53 | 0.52/0.59 | 0.57/0.65 | 0.51/0.50 | 0.54/0.46 | 0.45/0.49 | 0.58/0.58 | 0.58/0.58 | 0.44/0.53 | 0.44/0.53 | 0.52/0.48 |
| VMHC | 0.24/0.27 | 0.15/0.46 | 0.20/0.27 | 0.25/0.32 | 0.30/0.23 | 0.16/0.26 | 0.22/0.24 | 0.08/0.15 | 0.20/0.33 | 0.29/0.24 | 0.26/0.25 |

